# Plasticity and interactions in the odor responses of maxillary palps neurons in *Aedes aegypti*

**DOI:** 10.1101/2022.10.02.510498

**Authors:** Swikriti Saran Singh, Sanket Garg, Pranjul Singh, Smith Gupta, Abhinav Airan, Shefali Goyal, Nitin Gupta

## Abstract

Maxillary palps, in addition to the antennae, are major olfactory organs in mosquitoes and play an important role in the detection of human hosts. The sensory neurons of the maxillary palps reside in the capitate peg sensilla, each of which contains three neurons. In *Aedes aegypti*, the neuron with the largest spike amplitude in the sensillum is known to detect carbon dioxide. However, the responses of the other two neurons and the functional consequences of the grouping of these neurons within sensilla are not well understood. Here we identify odorants that activate the other two neurons. We detect a short-term plasticity in the odor-evoked local field potential of the sensillum and show that it originates in the spiking responses of the smallest-amplitude neuron, even though all three neurons contribute to the local field potential. We also detect inhibitory interactions among these neurons within the sensillum. We further show that the plasticity and the lateral interactions are functionally important as they affect the responses of the downstream projection neurons in the antennal lobe.

## Introduction

The *Aedes aegypti* mosquito is an anthropophilic vector for diseases like dengue, zika fever, yellow fever, and chikungunya. Female mosquitoes use a variety of cues to find human hosts for blood-feeding. These cues includes carbon dioxide (CO_2_) and components of the skin odor such as 1-octen-3-ol, 6-methyl-5-hepten-2-one, nonanal, 2-butanone, and many more (Kellogg 1970; Bernier et al. 2000, 2015; Logan et al. 2010; Cook et al. 2011; Saratha and Mathew 2016). The odorants are detected by receptors present on primarily two sensory organs: the antennae, which have been the subject of many studies, and the relatively less-studied maxillary palps. However, given that CO_2_ and many components of human odor are detected on the maxillary palps (Kellogg 1970; Omer and Gillies 1971; Grant et al. 1995; Grant and O’Connell 2007; Tauxe et al. 2013), a comprehensive understanding of the maxillary palp neurons is important for determining the neurobiological basis of host attraction in mosquitoes.

The maxillary palps contain several small basiconic sensilla present on their surface (Roth and Willis 1952; McIver and Charlton 1970). These sensilla, 29-35 in number in female *Aedes* mosquitoes, are known as the capitate pegs (cp), and each capitate peg sensillum houses three sensory neurons (McIver 1972; Mclver 1982). These neurons have been traditionally named according to the sizes of their spikes: cpA neuron has the largest spikes, cpB has intermediate spikes, and cpC has the smallest spikes (de Bruyne et al. 1999). In *Aedes, Anopheles*, and *Culex* mosquitoes, CO_2_ is detected by cpA (Kellogg 1970; Grant et al. 1995; Lu et al. 2007; Syed and Leal 2007). The same neuron also responds to some skin odorants (Lu et al. 2007; Tauxe et al. 2013). In *Anopheles* and *Culex*, 1-octen-3-ol is known to be detected by cpB (Lu et al. 2007; Syed and Leal 2007). In *Aedes*, while it is clear that 1-octen-3-ol is not detected by cpA, which of the other two neurons detects it remains puzzling. Some studies on *Aedes* have argued that 1-octen-3-ol is detected by cpB (Cook et al. 2011; Majeed et al. 2016; Herre et al. 2022), while others have argued it is detected by cpC (Grant and O’Connell 1996; Grant and Dickens 2011; Bohbot et al. 2013). Some authors have simply referred to the 1-octen-3-ol-sensitive neuron as the “small-amplitude cell” (DeGennaro et al. 2013). One reason that this confusion persists in *Aedes* is that the odors detected by the third neuron have remained unknown (Herre et al. 2022). Here, we resolve this issue by identifying different odors detected by cpB and cpC in *Aedes*.

The organization of the maxillary palps into multiple capitate peg sensilla, each containing three neurons, is conserved across different species of mosquitoes (Mclver 1982). However, the functional implications of this organization remain poorly understood. It is possible that this organization allows different subsets of ecologically relevant odorants to be detected by neurons with different physiological properties. We found that while cpA and cpB respond similarly to repeated presentations of an odorant, cpC responses are reduced with repeated exposure, providing a possible mechanism to prioritize the initial encounters with specific odors. The compartmental organization also creates room for lateral interactions between the sensory neurons (Vermeulen and Rospars 2004; Su et al. 2012; Miriyala et al. 2018; Zhang et al. 2019). In *Aedes*, we found that cpC activation inhibits cpA and cpB activity, and that these interactions affect the downstream responses in the antennal lobe. Together, our results reveal the responses, properties, and interactions of the maxillary palps neurons in *Aedes aegypti* and create a foundation for understanding the odor-processing underlying their attraction to human hosts.

## Results

### Odor responses of maxillary palp neurons

We performed *in vivo* single sensillum recordings (SSR) in capitate peg sensilla of female *Aedes aegypti* using glass pipettes. We saved the measurements with minimal filtering in the 0-10 KHz band to allow the detection of spikes and the local field potential (LFP). Precisely timed odors pulses, of typically 1-second duration in a 10-second trial, were delivered using a custom-built olfactometer (see **Methods**). Using custom routines for post-processing and applying thresholds on the second derivative of the recordings (see **Methods**), we observed three spike amplitudes, and accordingly labeled spikes as cpA, cpB, or cpC in the decreasing order of the amplitudes (**Figure 1a**).

**Figure 1:**
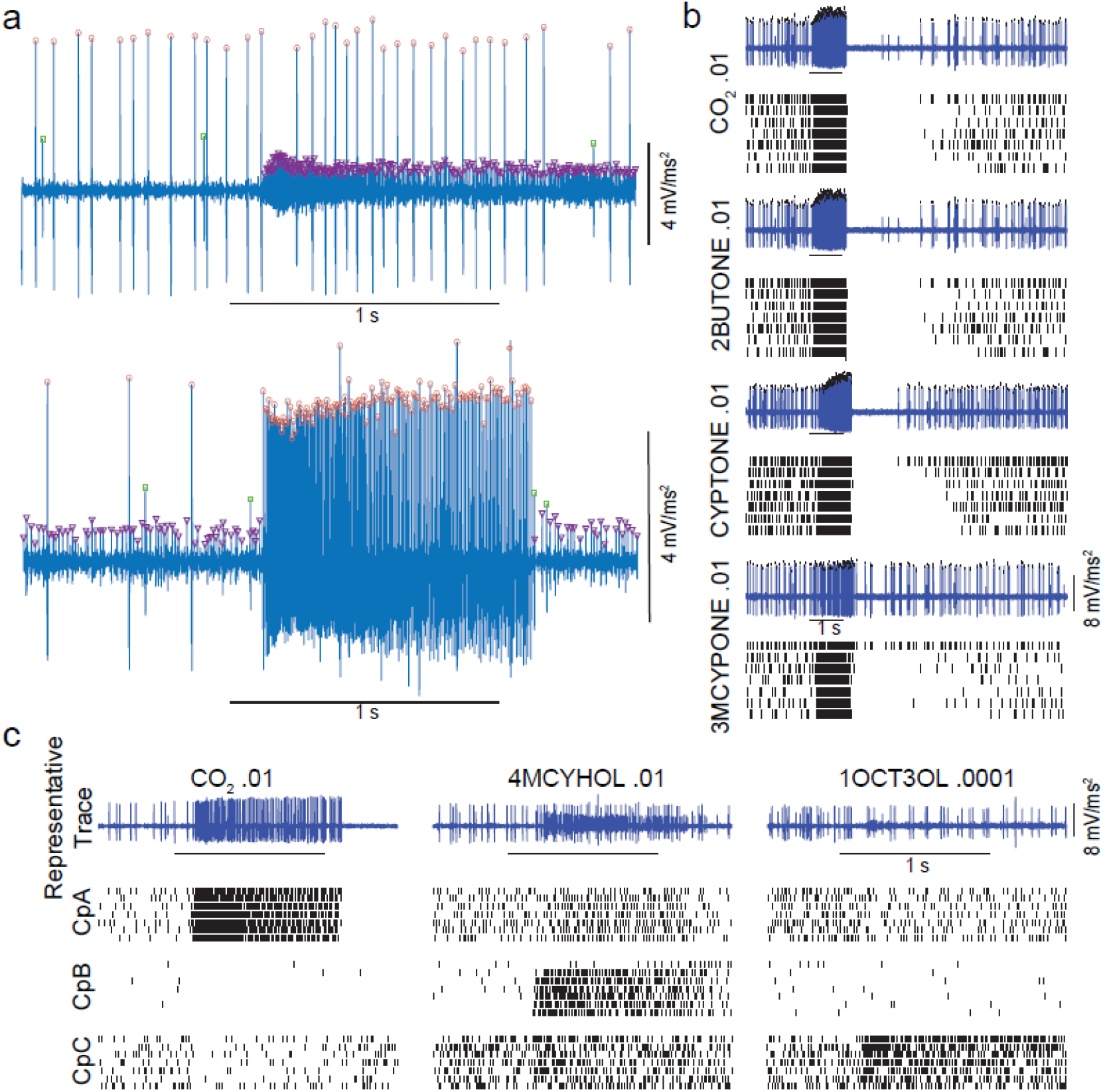
Single-sensillum recordings from capitate peg sensilla of A. aegypti. **a** A representative single-sensillum recording from a capitate peg sensillum in response to 1 s pulse (horizontal black lines) of 1-octen-3-ol (upper panel) and CO_2_ (lower panel). The traces shown are the second derivative of the raw signal. Individual spikes are labeled cpA (red circles), cpB (green squares), and cpC (purple triangles) based on their sizes. **b** Representative traces and rasters showing cpA responses to 1 s pulses of CO_2_, 2-butanone, cyclopentanone, and 3-methyl cyclopentanone. **c** Representative traces and rasters of cpA, cpB and cpC (from the same SSR) to 1 s pulses of CO_2_, 4-methylcyclohexanol, and 1-octen-3-ol show that cpA is activated by CO_2_, cpB by 4-methylcyclohexanol, and cpC by 1-octen-3-ol.

We first checked responses to CO_2_ and other known activators of cpA. Consistent with previous reports, we observed that cpA responded strongly to CO_2_, and also to some other odorants, such as cyclopentanone, 2-butanone, and 3-methyl cyclopentanone (**Figure 1b**). Next, we checked the responses to 1-octen-3-ol and found that it activates the neuron with the smallest spikes, cpC (**Figure 1c**). We also found that the remaining neuron with intermediate spikes, cpB, responded strongly to 4-methylcyclohexanol; Figure (**Figure 1c**) shows the identified spikes from the three neurons from the same SSR over multiple odor trials, confirming that cpA is activated by CO_2_, cpB by 4-methylcyclohexanol, and cpC by 1-octen-3-ol. In addition to activating cpB, 4-methylcyclohexanol also caused relatively weaker activation of cpA and cpC.

In recordings where cpA spikes were abundant, their large sizes made it technically difficult to detect any cpB or cpC spikes occurring in close temporal proximity. Thus, for odors that activated cpA, responses of cpB and cpC could not be measured reliably (**Figure 1a**). To overcome this limitation, we used Gr3-mutant mosquitoes (McMeniman et al. 2014). These mosquitoes have a deletion of 4 nucleotides in the Gr3 gene, and thus lack a functional copy of the Gr3 receptor, which is essential for CO_2_ detection by the cpA neuron (Erdelyan et al. 2012; McMeniman et al. 2014). Consistent with the earlier results (McMeniman et al. 2014), we found that Gr3 mutants did not show spontaneous or CO_2_-evoked cpA spikes (**Figure 2a**). This allowed us to reliably observe the responses of cpB and cpC neurons to different odors. Recordings from these mosquitoes further confirmed that 1-octen-3-ol activates only cpC, while 4-methylcyclohexanol activates cpB strongly and cpC weakly. We also attempted to identify other odors that activate cpB. In our test panel, cpB responded to fewer odors than cpC (**Figure 2a**). Overall, we examined cpB responses to 16 odorants (some at multiple concentrations) and 2 solvents (mineral oil and water) in multiple SSR experiments performed on Gr3 mutants, and found that cpB consistently showed the strongest response to 4-methylcyclohexanol and also showed weaker responses to 1-hexen-3-ol, (S)-limonene, 3-methyl-cyclopentanone and dimethyl trisulfide (**Figure 2b**).

**Figure 2:**
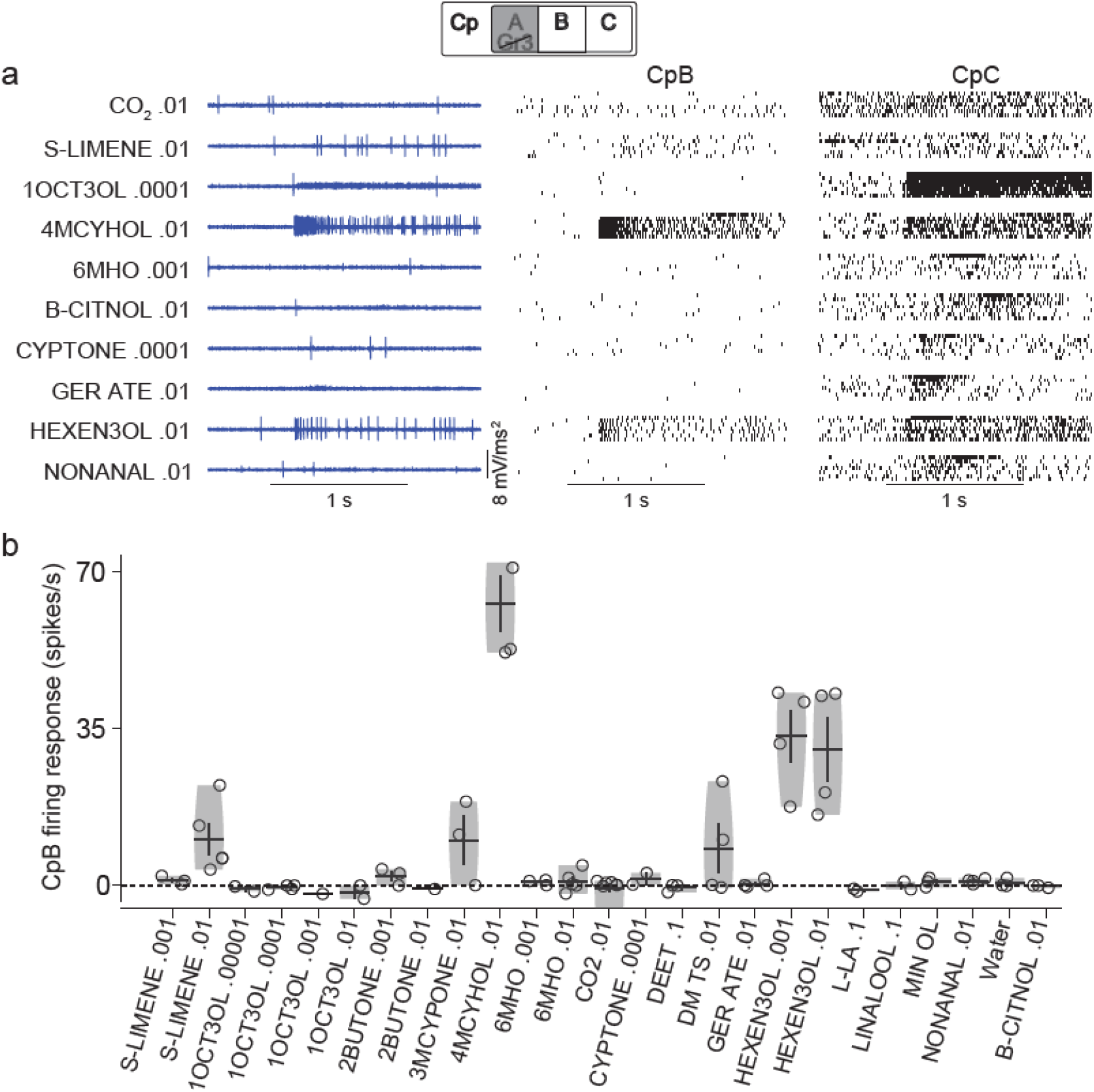
Odors that activate cpB neurons. **a** Representative traces and rasters of cpB and cpC in response to 1 s pulses of different odors in Gr3 mutants. **b** Violin plots showing odor-evoked responses of cpB for different odors. Each point represents the odor-evoked cpB response from one sensillum (averaged over 7 trials for each odor). For odor acronyms, see Methods.

### Plasticity in LFP responses

Our SSR recordings from the capitate peg sensillum, obtained without high-pass filtering of the signal, showed that the odor-elicited spikes were accompanied by deflections in the LFP. CO_2_, for example, showed a sharp downward deflection in the LFP along with the increase in cpA spikes (**Figure 3a**). In our experiments, each odor stimulus of 1 s duration was presented typically 7 times with a gap of 10 s between the presentations. The LFP responses generated by CO_2_ were similar across the 7 trials (**Figure 3b**).

**Figure 3:**
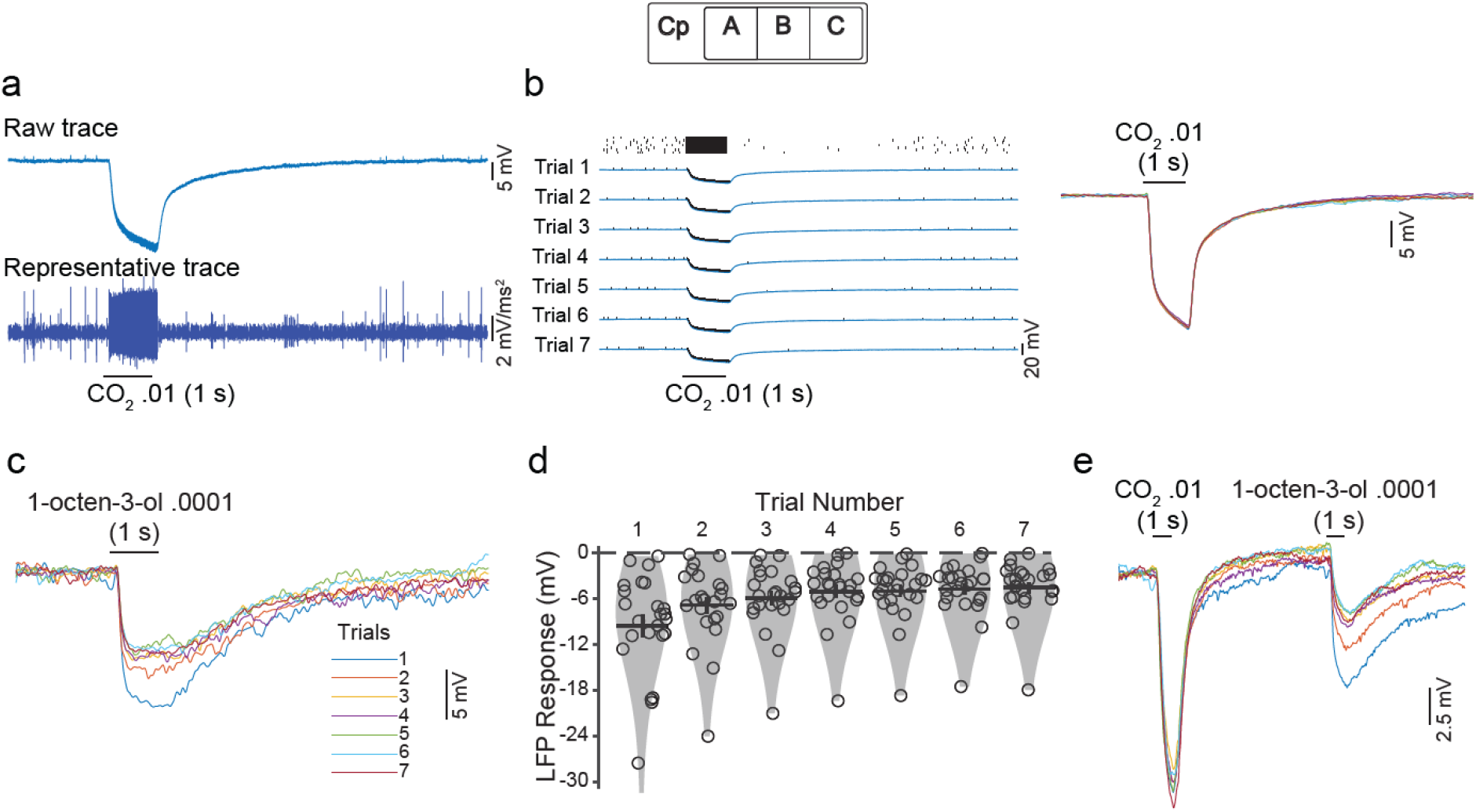
Plasticity in the LFP response to 1-octen-3-ol. **a** A representative SSR shows the downward deflection in the LFP (upper trace) and an increase in cpA spiking activity (lower trace) in response to a 1 s pulse of CO_2_. **b** Left: raster plots and recording traces from 7 trials showing cpA spikes along with the downward deflections. Right: Recording traces from the 7 trials of CO_2_, placed over each other, show that the responses are very similar in all the trials. **c** Representative traces showing reduction in LFP responses to 1-octen-3-ol over 7 trials. Average background activity (pre-odor) was subtracted from each trial to equalize the baselines and highlight the odor-evoked responses. **d** Recordings of 24 different sensilla with 1-octen-3-ol 0.0001 show stronger LFP deflections in the initial trials. **e** Representative traces showing LFP response plasticity with 1-octen-3-ol but not with CO_2_ even when the two odors were delivered alternately.

Surprisingly, we observed a different pattern with 1-octen-3-ol: LFP response was the largest in the first trial and reduced with subsequent trials (**Figure 3c**). Over a dataset of 24 recordings from 20 mosquitoes tested with 1-octen-3-ol 0.0001 (**Figure 3d**), the LFP response was significantly larger in magnitude in the first trial than the second trial (P = 2.7 x 10^-5^, n = 24, sign-rank test), in the second trial than the third trial (P = 6.6 x 10^-3^), and in the third trial than the fourth trial (P = 1.5 x 10^-3^). Over all trials, the magnitude of the LFP response showed a significant negative correlation with the trial number (R = 0.87, P = 0.01, n = 7 trials). We performed another experiment in which we delivered alternate pulses of CO_2_ and 1-octen-3-ol over 7 trials. Here too, we found that the LFP response to CO_2_ remained stable while the LFP response to 1-octen-3-ol reduced over the trials (**Figure 3e**).

To check if the reduction in LFP response to 1-octen-3-ol was caused by a gradual reduction in the amount of odor being delivered to the animal, we measured the odor delivery profile using a photo-ionization detector (PID). We found that the amount of odor delivered did not reduce over the trials (**Figure S1a**); on the contrary, the amount of 1-octen-3-ol delivered was typically lower in the first trial than the subsequent trials (as replacing the background air with the odorized air in the odor delivery tubes took some time). Similar observations were made on six different days (**Figure S1b**). Thus, the reduction in LFP response over trials could not be explained by a reduction in the amount of odor delivered. Rather, these results provide evidence for a plasticity in the LFP response to 1-octen-3-ol in the capitate peg sensillum.

### Mechanism of the plasticity in LFP responses

We checked the effect of odor concentration and odor pulse duration on the plasticity. The above results were obtained with 0.0001 concentration of 1-octen-3-ol. When we checked the responses for 10-fold lower (0.00001), 10-folder higher (0.001) and 100-fold higher (0.01) concentrations, we still found the plasticity in the LFP responses (**Figure S2a**). When we reduced the pulse duration from 1 s to 0.5 s for 0.0001 and 0.00001 concentrations of 1-octen-3-ol, the plasticity was still observed (**Figure S2b**). When we further reduced the pulse duration to only 0.25 s, we observed the plasticity for 0.0001 concentration but did not observe it for 0.00001 concentration (**Figure S2c**). Thus, except for the lowest pulse width at the lowest concentration, the plasticity was observed for all combinations of concentrations and pulse widths that we tested.

We next asked if the observed plasticity in the LFP response exists for multiple odors or is unique to 1-octen-3-ol. We checked the LFP responses across trials for 16 different odors (**Figure S3**). We found that all of these odors generated LFP deflections in the capitate peg sensillum, but the reduction in the LFP response over trials was observed for some of these odors, such as (S)-limonene, β-citronellol, and geranyl acetate. Thus, the plasticity of the LFP response is odor specific. For some odors, such as linalool, 4-methylcyclohexanol, cyclopentanone and 3-methyl-cyclopentanone, an opposite trend was observed: the LFP deflection was weaker in the first trial compared to later trials, which is likely explained by lower odor delivery in the first trial.

We next sought to understand the involvement of cpA, cpB, and cpC neurons in the plasticity. Experiments in *Drosophila* suggest that LFP deflections arise due to transduction currents in the receptor neurons (Nagel and Wilson 2011) and different neurons within a sensillum can contribute differently to the LFP (Zhang et al. 2019). We first checked whether all three neurons in the capitate peg sensillum contribute to the LFP.

While cpA primarily expresses gustatory receptors for the detection of CO_2_, the other two neurons express odorant receptors (ORs), along with some ionotropic receptors (IRs) (Bohbot et al. 2014; Herre et al. 2022). To check the contribution of cpA firing to LFP, we performed experiments in Orco-mutant mosquitoes, which lack the functional olfactory receptor co-receptor (Orco) required for the activity of all ORs (DeGennaro et al. 2013). Thus, Orco-mutant mosquitoes have greatly reduced cpB and cpC responses, enabling us to check the relationship between cpA firing and LFP more clearly. We compared cpA firing rate and LFP deflection in 200-ms bins over a 4-s period starting from the onset of odor delivery and checked the correlation between the two quantities in each recording (**Figure S4**). A negative correlation indicates that bins with high cpA firing had more negative LFP deflections. Across multiple recordings with odors that activated cpA, we found the mean correlation to be significantly below zero (R = −0.71 ± 0.02, P = 8.29 x 10^-6^, n = 26 recordings, sign-rank test; **Figure 4a**). This result suggests that cpA firing contributes to the LFP. Next, we used Gr3-mutant mosquitoes to clearly identify cpB and cpC spikes and check their contribution to the LFP. Again, we took multiple recordings in which cpB was activated by the odor, and for each recording calculated the correlation between cpB firing rate and LFP over different temporal bins. A negative correlation (R = −0.90 ± 0.01, P = 1.32 x 10^-4^, n = 19 recordings, sign-rank test; **Figure 4b**) suggests that cpB firing contributes to the LFP. Finally, a similar analysis for cpC neurons also showed a negative correlation (R = −0.81 ± 0.02, P = 8.29 x 10^-6^, n = 26 recordings, sign-rank test; **Figure 4c**). Thus, spikes of all three neurons contribute to the LFP in the *Aedes* capitate peg sensillum.

**Figure 4:**
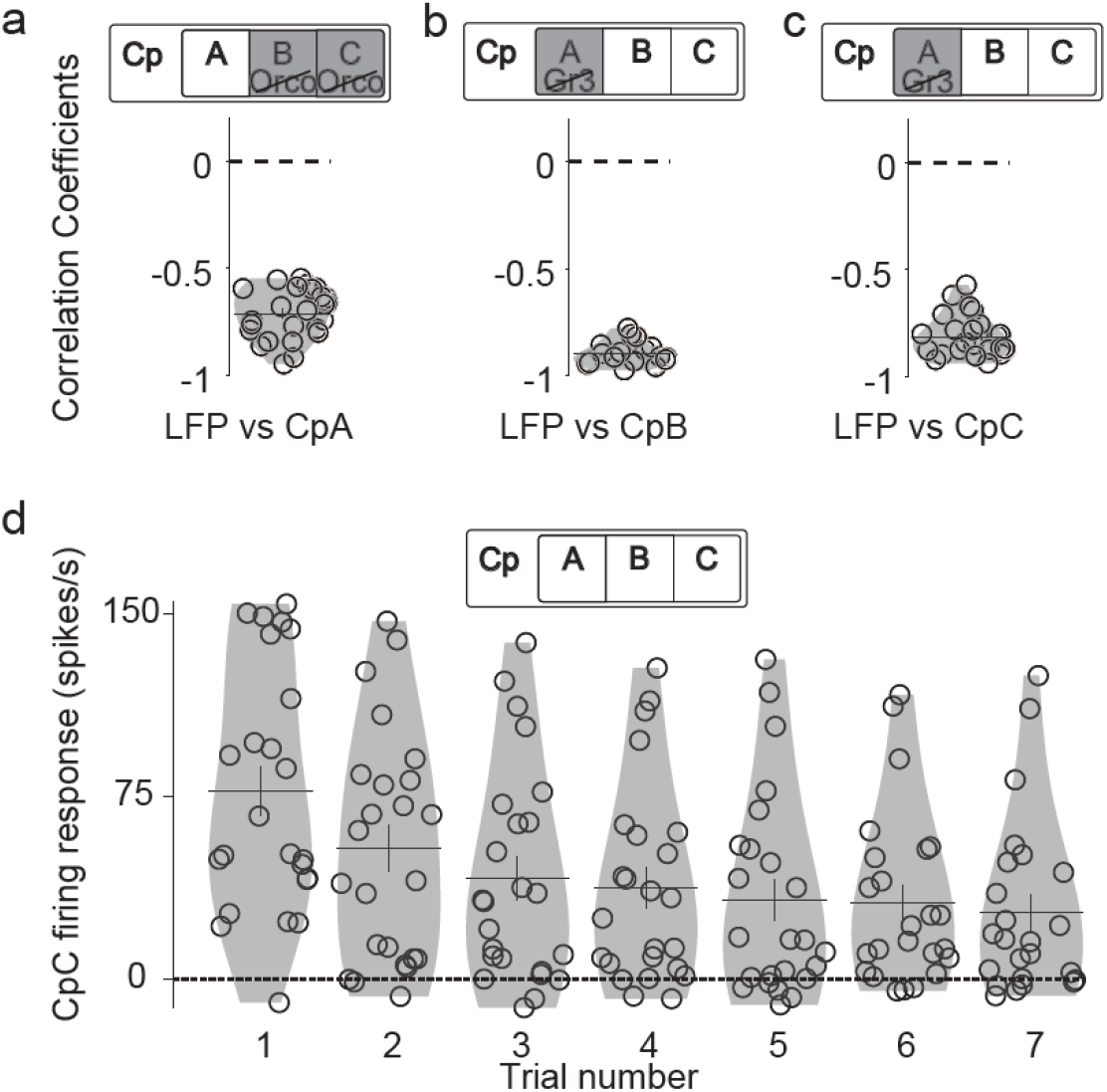
All three neurons in the capitate peg sensillum contribute to LFP. **a** Correlations between LFP responses and cpA firing responses from Orco-mutant mosquitoes (n = 26 SSRs). **b** Correlations between LFP responses and cpB firing responses from Gr3-mutant mosquitoes (n = 19 SSRs). **c** Correlations between LFP responses and cpC firing responses from Gr3-mutant mosquitoes (n = 26 SSRs). Note that the correlation values are negative in all cases. **d** Violin plots show a decrease in cpC firing rates in response to 1-octen-3-ol 0.0001 over trials.

As all three neurons contribute to the LFP, it is not immediately obvious if all of them are also involved in the LFP plasticity. Since we observed the plasticity with 1-octen-3-ol 0.0001, which activates cpC, we expected it might be caused by a corresponding decrease in the odor-evoked firing response of cpC over trials. Indeed, we found that the change in cpC firing evoked by 1-octen-3-ol 0.0001 was significantly smaller in the second trial compared to the first trial (P = 6.06 x 10^-4^; n = 24 recordings; sign-rank test), and continued to decrease over the subsequent trials (**Figure 4d**). Thus, the plasticity in the LFP response to 1-octen-3-ol can be explained by the plasticity in the cpC firing. In contrast, since we did not observe LFP plasticity with CO_2_, which is detected by cpA, we expected cpA firing to also lack the plasticity. **Figure S5a** shows LFP and cpA firing responses for a few odors in the same sensillum in Orco-mutant mosquitoes: no reduction is seen over trials (the increase from the first to the second trial in some odors is due to increased odor delivery, as discussed above). The same trend of no reduction in cpA firing over trials was seen across recordings from multiple sensilla (**Figure S5b**). Lastly, we checked if cpB firing shows plasticity over trials. Using Gr3-mutant mosquitoes and odors that activate cpB, we did not find evidence for plasticity in cpB firing (**Figure S6**). In summary, only cpC shows plasticity in its firing rate over trials, and that appears to be responsible for the plasticity in the LFP response.

If cpC firing plasticity is indeed responsible for the LFP plasticity, odors that show more plasticity in the cpC firing should also result in more plasticity in the LFP responses. To test this hypothesis, we checked responses to multiple odors in Gr3 mutants. In addition to 1-octen-3-ol, other tested odors also activated cpC to different extents and exhibited plasticity in the cpC firing rate and in the LFP responses (**Figure 5a, b, c**). We quantified the level of LFP plasticity in each recording by fitting a line to the negative peak values for the 7 trials and calculating its slope (**Figure 5d**). A positive slope for LFP indicates that the LFP response became less negative (i.e., weaker) over the trials. The level of plasticity in cpC firing rate for an odor was quantified by fitting a line to the cpC firing rates across the 7 trials and calculating its slope (**Figure 5e**). A negative value of the slope for cpC indicates that the cpC firing rate reduced over the trials. We checked the correlation between the two slopes from multiple odor recordings from 5 different SSRs, and found that more positive slopes for LFP response were indeed associated with more negative slopes for cpC firing (R = −0.65, P = 5.37 x 10^-6^; n = 40 recordings; **Figure 5f**). Thus, recordings that had more plasticity in cpC firing also had more plasticity in the LFP response.

**Figure 5:**
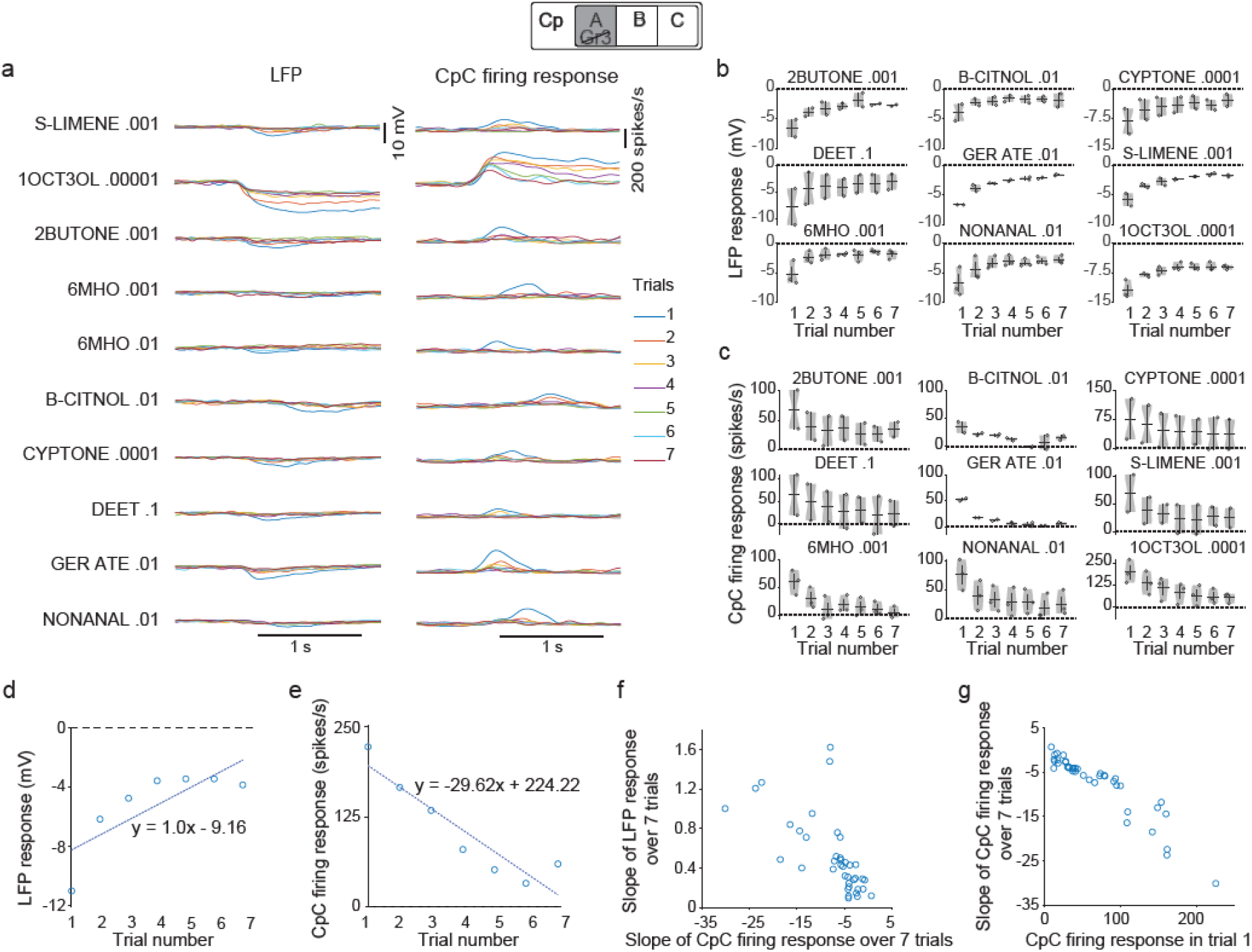
Plasticity in LFP is related to the plasticity in cpC firing. **a** LFP and cpC firing rates from the same sensillum in a Gr3-mutant mosquito in response to (S)-limonene (S-LIMENE), 1-octen-3-ol (1OCT3OL), 2-butanone (2BUTONE), 6-methyl-5-hepten-2-one (6MHO), β-citronellol (B-CITNOL), cyclopentanone (CYPTONE), DEET, geranyl acetate (GER ATE), and nonanal. **b** Violin plots showing data from multiple recordings confirm plasticity in LFP responses over trials. **c** Violin plots show plasticity in cpC firing rate responses. **d** Scatter plots between LFP responses and trial numbers in a recording from a Gr3 mutant. A linear fit was obtained and the slope of the fitted line indicated the level of plasticity (1.01 in this example). A positive value of the slope indicates the expected plasticity. **e** Change in cpC firing responses versus trial number. A negative value of the slope of the fitted line indicates the expected plasticity (−29.62 in this example). **f** Plasticity in LFP was correlated with the plasticity in cpC firing rate. **g** Plasticity in cpC firing rate was correlated with the firing rate of the cpC neuron in the first trial.

What determines the level of plasticity in cpC firing rate? We checked if odors that activate cpC neuron more strongly also result in more plasticity in the cpC firing rate. We calculated the correlation between the slope of the firing rate and the firing rate itself (in the first trial), and found a strong negative correlation between them (R = −0.93, P = 4.43 x 10^-18^; n = 40 recordings; **Figure 5g**). Thus, we conclude that odors that activate the cpC neuron more strongly also result in more plasticity in the cpC firing rate over trials, which in turn results in more plasticity in the LFP response.

Next, we considered other possible factors that may contribute to the plasticity in the LFP response. In our experiments so far, a trial was of 10 s duration, and the odor pulse was delivered in the 2-3 s interval. Although the LFP response started with the onset of the odor pulse and started returning towards the baseline at the end of the 1 s pulse, in some cases the return was slow and the LFP did not reach the baseline by the end of the trial (**Figure 3c**). In such cases, the next trial would start from a slightly different baseline, and since we calculate the LFP response as the difference between the peak response and the baseline, this may potentially contribute to the appearance of a plasticity effect. However, this is unlikely to be the major reason for the plasticity we observed, for two reasons. First, odors for which the LFP returned to the baseline completely by the end of each trial, such as nonanal and 6-methyl-5-hepten-2-one (**Figure S7a**), also showed plasticity in the LFP response (**Figure 5b**). Second, even when we increased the trial duration to 20 s, the plasticity was observed (**Figure S7b**). Taken together, our results suggest that the plasticity in the LFP derives primarily from the plasticity in the cpC firing rate, which appears to be a physiological property of the cpC neuron. While most sensory neurons show adaptation during a continued stimulus presentation, the plasticity we observed was unique to cpC and lasted for at least 20 s after the stimulus.

### Significance of the cpC plasticity to downstream neurons

One way to assess the functional significance of the plasticity observed in cpC responses is to check if it affects the responses of neurons at the next layer of information processing. Maxillary palps neurons send their axonal projections to three glomeruli in the antennal lobe, I-MD1, I-MD2, and I-MD3 (Ignell et al. 2005), where they synapse with projection neurons (PNs). Genetic labeling experiments have shown that cpA neurons project to I-MD1 glomerulus, while cpB and cpC project to the other two glomeruli (Herre et al. 2022). Using patch-clamp recordings and post-hoc dye filling, we were able to observe the odor responses of some PNs innervating these glomeruli (Singh et al. 2022). As we show below, I-MD2 PN responded strongly to 4-methylcyclohexanol but not to CO_2_ or 1-octen-3-ol, while I-MD3 PNs responded to 1-octen-3-ol: thus, we inferred that cpB neurons project to I-MD2 and cpC neurons project to I-MD3 glomerulus.

We checked if the I-MD3 PNs, downstream to cpC neurons, also show plasticity in their responses over trials. **Figures 6a-d** shows the responses of an I-MD3 PN to CO_2_ and to three different concentrations of 1-octen-3-ol. This PN did not respond to CO_2_ but responded with increased firing rate to 1-octen-3- ol, with strong responses to 0.01 and 0.001 concentrations and moderate response to 0.0001 concentration. Moreover, for the higher concentrations, the response was stronger in the first trial and reduced over trials, showing that the plasticity is also present downstream to cpC (**Figure 6b, c, d**). We calculated the slope of the fit to the firing rate over trials to quantify the plasticity in multiple PNs tested with the same odor and found that I-MD3 PNs showed the effect (mean slope = −3.79 ± 1.28; n = 9 PNs; P = 7.81 x 10^-3^; sign-rank test; **Figure 6e**), while most of the other glomeruli did not show this effect (**Figure S8**). Many of the PNs tested here receive input from antennal sensilla, and the lack of the plasticity in their responses is consistent with the idea that the observed plasticity may be a property of cpC neurons on the maxillary palp.

**Figure 6:**
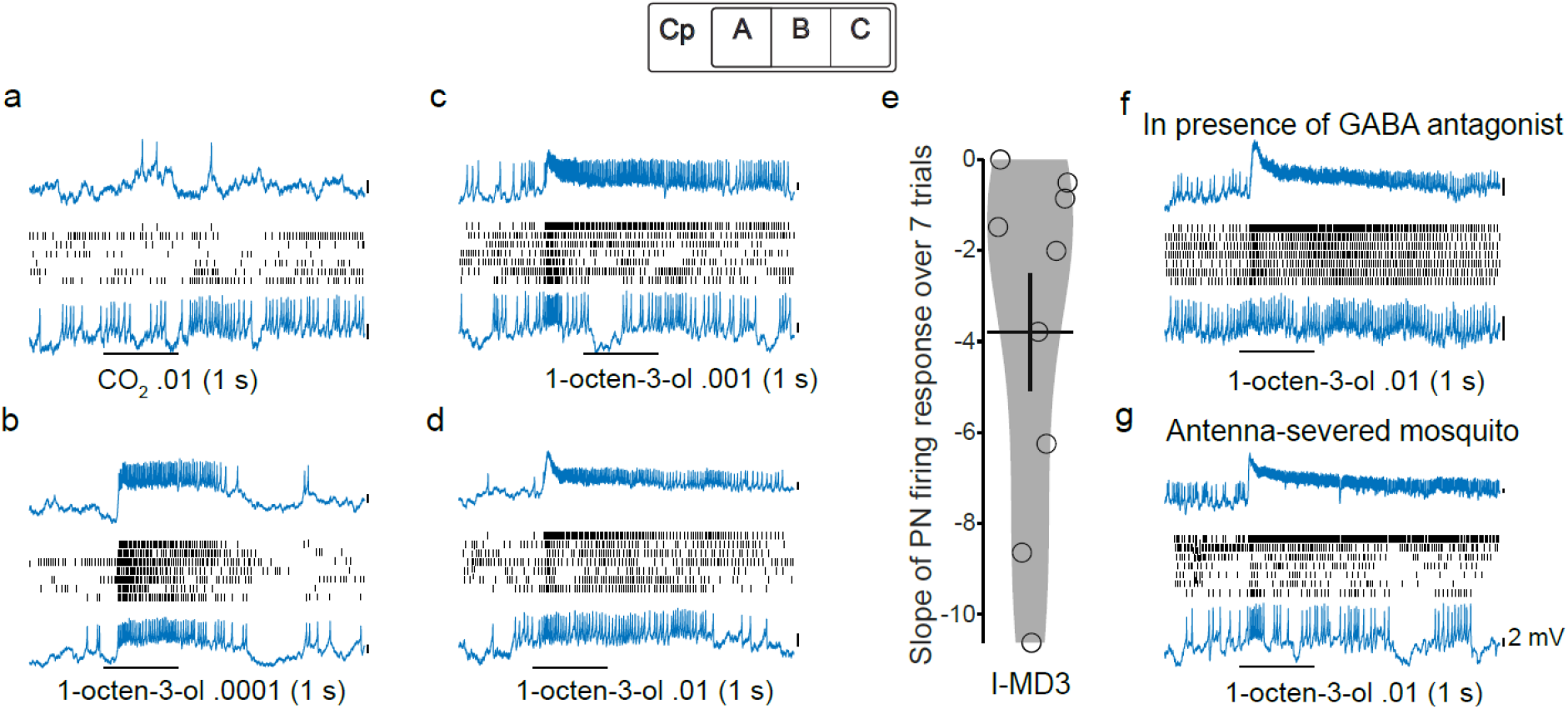
Plasticity in a projection neuron that receives input from cpC. **a-d** Representative traces (first and last trials on top and bottom, respectively) and rasters of an I-MD3 PN’s responses to CO_2_ .01 (**a**), 1-octen-3-ol .0001 (**b**), 1-octen-3-ol .001 (**c**), and 1-octen-3ol .01 (**d**). **e** Violin plot showing plasticity values in I-MD3 PNs (n = 9) in response to 1-octen-3-ol .01. Each point on represents the slope of the fitted line for PN firing responses across the 7 trials; negative slope indicates plasticity. **f** Representative traces and rasters of an I-MD3 PN in response to 1-octen-3-ol .01 show plasticity in the presence of a GABA antagonist (picrotoxin 10 μM). This is the same cell as shown in **a-d**. **g** Recording from an I-MD3 PN in a mosquito in which antennae were cut before the experiment. Note the strongest response in the first trial. Vertical scale bars in all panels, 2 mV.

The insect antennal lobe involves extensive lateral interactions mediated by local neurons (Chou et al. 2010). Thus, the activity of a PN depends not only on the sensory input it receives from the sensory neurons, but also on the lateral inputs received from other neurons within the antennal lobe. Gamma-aminobutyric acid (GABA) is one of the primary inhibitory neurotransmitters inside the *Aedes* brain (Matthews et al. 2016), and GABA-positive LNs have been described in the antennal lobe (Singh et al. 2022). Could lateral interactions also contribute to the observed plasticity in I-MD3 PNs? Two observations did not support this idea. First, for the same I-MD3 PN shown in **Figure 6d**, we measured the response to 1-octen-3-ol 0.01 after application of picrotoxin, a GABA-A receptor antagonist, and found that the plasticity was still present **Figure 6f**). Second, we measured the activity of an I-MD3 PN in another experimental preparation in which the antennae were severed. In this preparation, the inputs to I-MD3 from the palp sensory neurons were intact, but the lateral inputs from other neurons in the antennal lobe that are normally activated by the sensory neurons on the antenna would have been curtailed. Even in this preparation, we observed the same plasticity in the I-MD3 PN (**Figure 6g**). Together, these results show that the plasticity in the 1-octen-3-ol response of I-MD3 PN is unlikely to be mediated by lateral interactions and derives primarily from cpC sensory neurons.

### Inhibitory interactions within the capitate peg sensillum

While measuring the responses to 1-octen-3-ol in the capitate peg sensillum, we observed a curious phenomenon. The odor activated the cpC neuron as expected, but in many recordings, we also observed a reduction in the spontaneous firing of the cpA neuron at the same time (**Figure 7a**). By pooling data from multiple recordings, we confirmed that stimulation with 1-octen-3-ol 0.01 resulted in a significant reduction in the spontaneous firing rate of cpA (mean change in firing rate: −20.22 ± 2.53 spikes/s, P = 5.25 x 10^-6^; sign-rank test; n = 28 recordings; **Figure 7b**). As cpA spikes are much larger than cpC spikes, the inhibitory effect cannot be explained by incomplete detection of cpA spikes when cpC is firing at high rate.

**Figure 7:**
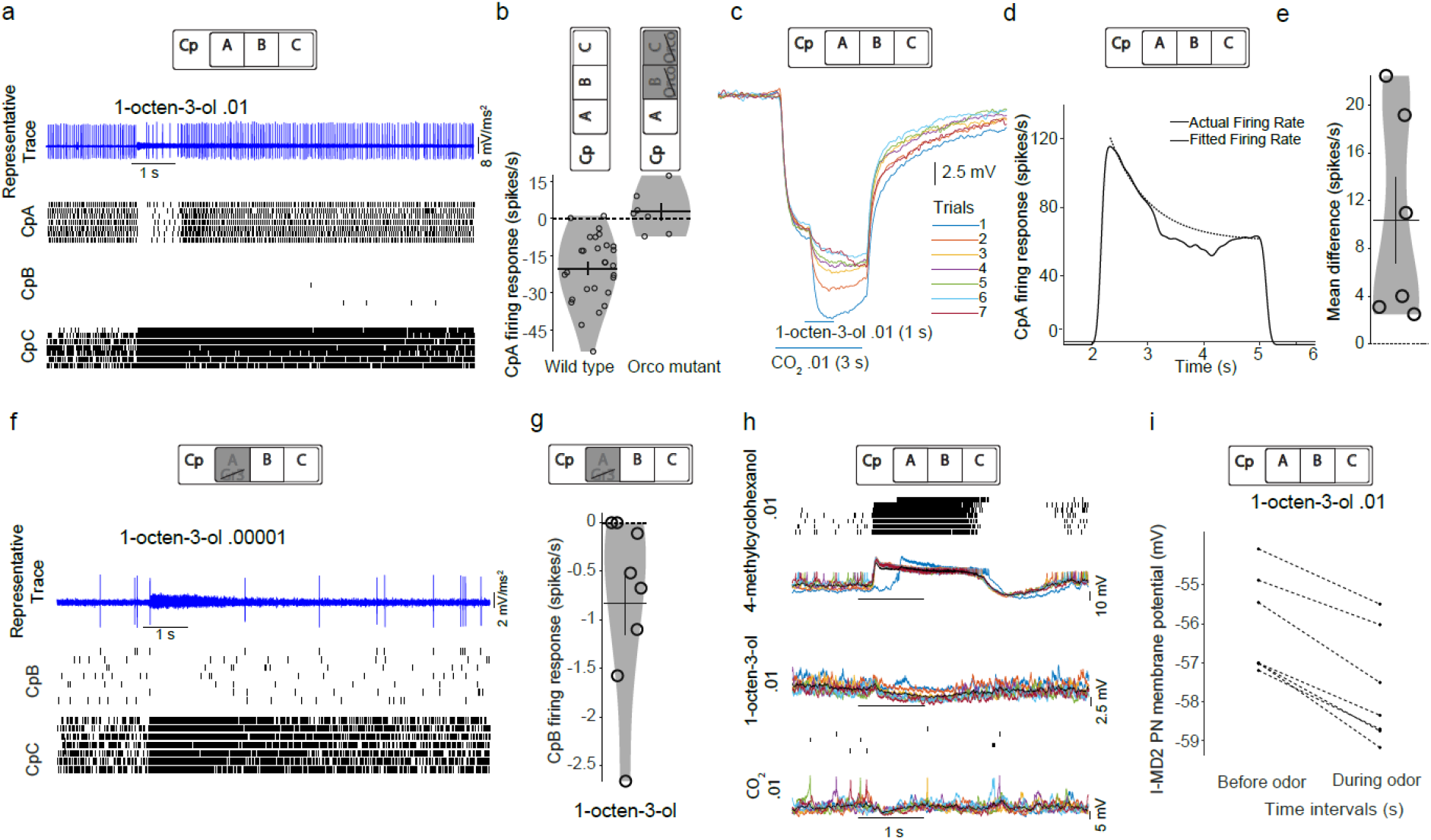
Inhibition in cpA and cpB neurons during cpC activation. **a** Representative trace and rasters of cpA, cpB, and cpC in response to 1 s pulse of 1-octen-3-ol .01 in a 10 s trial show an increase in cpC firing and a concurrent inhibition in cpA spontaneous firing; cpB did not show much activity in this recording. **b** Violin plots show inhibition in the firing of cpA in response to 1-octen-3-ol .01 in wild-type (left, n = 28 SSRs) but not in orco-mutant (right, n = 7 SSRs) mosquitoes. **c** Representative traces showing LFP responses to an overlapping pulse of 1-octen-3ol .01 (1s) in the middle of a prolonged CO_2_ .01 pulse (3 s). **d** Dotted line shows an exponential decay curve fitted to cpA firing rate (solid line) with stimulus pattern mentioned in **c** (see Methods). **e** Each point represents mean difference (fitted – actual) from 6 mosquitoes; a positive value indicates inhibition of CO_2_-evoked cpA firing response when 1-octen-3-ol is given on top of it. **f** Representative traces and rasters of cpB and cpC in response to 1 s pulse of 1-octen-3-ol .01 from Gr3-mutant mosquitoes show an increase in cpC firing and a concurrent inhibition in cpB spontaneous firing. **g** Violin plot shows a reduction in cpB firing upon 1-octen-3-ol stimulation in 8 Gr3-mutant recordings. **h** Rasters and representative traces of I-MD2 PN in response to 4-methylcyclohexanol .01, 1-octen-3-ol .01, and CO_2_ .01. Each panel displays rasters and membrane potentials from 7 trials (color code same as **c**). **i** Comparing average membrane potential before odor pulse and during odor pulse in 1-octen-3-ol trials. A decrease in I-MD2 PN membrane potential during odor pulse indicates a reduction in the synaptic input during 1-octen-3ol stimulation (P = 0.016, n = 7 trials; sign-rank test).

The reduction in cpA firing may be caused by direct interaction of 1-octen-3-ol with a receptor on cpA that has an inhibitory effect, or it may be caused by lateral inhibition from cpC activated by 1-octen-3-ol. If the former is true, we should observe the inhibition in cpA with 1-octen-3-ol even when we block the activation of cpC. We tested this hypothesis using recordings from Orco-mutant mosquitoes, in which cpC lacks the functional OR complex required to detect 1-octen-3-ol. Note that cpA expresses only GRs and some IRs (Herre et al. 2022), and is therefore not directly affected by the Orco mutation.

We found that the inhibition of cpA firing upon 1-octen-3-ol stimulation was abolished in Orco mutants (mean change in firing rate: 2.67 ± 3.21 spikes/s, P = 0.47, sign-rank test, n = 7 recordings; **Figure 7b**). Therefore, we conclude that the reduction in cpA firing is caused by lateral inhibition from cpC to cpA, possibly via ephaptic coupling (Su et al. 2012).

The above experiments showed that activation of cpC can inhibit the spontaneous firing of cpA. We next asked whether activation of cpC also inhibits odor-evoked firing in cpA. To check this, we performed experiments in which a 3 s pulse of CO_2_ was given in the 2-5 s interval within a 20 s trial, and on top of it a 1 s pulse of 1-octen-3-ol was given in the 3-4 s interval. The long pulse of CO_2_ activated cpA and resulted in a strong LFP response, which was further strengthened by the pulse of 1-octen-3-ol, suggesting that overall firing in the sensillum increased when 1-octen-3-ol was delivered (**Figure 7c**). Because of adaptation, the instantaneous firing rate of cpA gradually reduced within the 3 s CO_2_ pulse. However, a careful look at the temporal profile of the cpA firing rate suggested an additional reduction in cpA firing when 1-octen-3-ol was delivered (**Figure 7d**). To quantify this additional inhibition caused by 1-octen-3-ol, we fitted a curve to the cpA firing rate in the periods before and after the 1-octen-3-ol pulse, and then calculated the difference between the fitted and the actual firing rate (**see Methods**). The fit approximates the firing rate profile expected in the absence of 1-octen-3-ol pulse. We found that the actual firing rate was on average lower than the fit by −10.37 ± 3.56 spikes/s (P = 0.03; sign-rank test; n = 6 recordings; **Figure 7e**). Thus, we conclude that the lateral inhibition not only affects cpA’s spontaneous firing but also affects its odor-evoked firing.

Neuron cpC ephaptically inhibits neuron cpA spontaneous response; we asked whether it also inhibits neuron cpB’s spontaneous response. We used Gr-3 mutant mosquitoes to check this, where neuron cpB and cpC responses are not affected by large spiking neuron cpA. Since 1-octen-3-ol activates cpC neurons, we asked whether 1-octen-3-ol activation inhibits cpB neuron response as it does with cpA neurons. **Figure 7f** shows when there is an activity in cpC neuron, there is inhibition in cpB neuron response. Recordings from multiple SSRs confirmed inhibition in cpB spontaneous firing response with an increase in cpC firing response during 1-octen-3-ol stimulation (mean change in cpB firing rate: −0.83 ± 0.33 spikes/s, P = 0.03; sign-rank test; n = 8 recordings; **Figure 7g**).

Next we asked if this lateral inhibition among the sensory neurons in the maxillary palps affects the downstream responses. We found that I-MD2 PN responds strongly to 4-methylcyclohexanol, but not to CO_2_ or 1-octen-3-ol (**Figure 7h**). Therefore, as discussed above, we inferred that it receives direct input from cpB neurons on the maxillary palp. Consistent with the lateral inhibition of cpB during 1-octen-3-ol stimulation, the membrane potential of the I-MD2 PN also reduced during 1-octen-3-ol stimulation (P = 0.016, n = 7 trials; sign-rank test; **Figure 7i**). This result is consistent with the idea that inhibitory interactions among the maxillary palps neurons affect the responses of downstream PNs in the antennal lobe.

## Discussion

In summary, our results revealed the odors detected by different neurons in the capitate peg sensillum on the maxillary palps of *Aedes aegypti*. Our LFP measurements from the sensillum showed evidence for plasticity over trials, which lasted for at least 20 s and was observed for different odor pulse lengths and different odor concentrations. We showed that while all three neurons in the sensillum contribute to the LFP, the plasticity in the LFP originates from a similar plasticity in the firing rate of only the cpC neuron. The firing was seen for most odors that activated cpC, and the magnitude of plasticity was proportional to the strength of cpC firing. Our recordings also revealed lateral inhibition within the capitate peg sensillum of Aedes aegypti: cpC spikes inhibited cpA and cpB spikes.

We identified and measured the responses of PNs in the antennal lobe that are downstream of the maxillary palp neurons and showed that both the plasticity and the lateral interactions observed in the sensory neurons affect the responses of the downstream PNs.

The capitate peg sensilla on the maxillary palps in all well-studied mosquito species are known to contain three neurons, of which cpA is sensitive to CO_2_ and one of the other two neurons to 1-octen-3-ol. However, the identity of the neuron that detects 1-octen-3-ol in *Aedes aegypti* has remained contentious, with some studies arguing for cpB (Cook et al. 2011; Majeed et al. 2016; Herre et al. 2022) and others for cpC (Grant and O’Connell 1996; Grant and Dickens 2011; Bohbot et al. 2013). Our results clearly show that 1-octen-3-ol is detected by cpC. If cpA and cpC spikes are present in a recording but cpB spikes are absent, then those cpC spikes will appear as the second largest spikes and may be incorrectly interpreted as cpB spikes (as the naming of the neurons is based on their *relative* spike sizes). This can happen because the odors that activate the remaining neuron (other than the CO_2_-sensitive neuron and the 1-octen-3-ol-sensitive neuron) were not known so far in *Aedes aegypti* (Herre et al. 2022); the spontaneous firing may also be absent in some cases as it varies with the experimental preparations. We speculate that a lack of actual cpB spikes in some studies likely explains why the 1-octen-3-ol-sensitive neuron was labeled cpB in those studies. Our results also show that cpB responds strongly to 4-methylcyclohexanol and weakly to others including 1-hexen-3-ol, (S)-limonene, 3-methyl-cyclopentanone and dimethyl trisulfide. This information should help future studies in separating the three neurons more reliably.

It is also conceivable that the exact position of the electrode in the sensillum can change the relative sizes of spikes from the two smaller-spiked neurons; however, in our recordings, we did not find any evidence for this: spikes in the 1-octen-3-ol-sensitive neuron were always smaller in size than in the 4-methylcyclohexanol-sensitive neuron. A recent study in *Drosophila* has shown that the spike size is related to the morphological size of the neuron in the sensillum (Zhang et al. 2019): this also suggests that the measured spike sizes should not be very sensitive to the placement of the electrode. Another possible reason for the difference in the studies reporting cpB or cpC as responding to 1-octen-3-ol could be the possible heterogeneity among different capitate peg sensilla on the maxillary palps (Herre et al. 2022). Even this heterogeneity is unlikely to result in a systematic bias among studies, as most studies like ours randomly select one of the many sensilla on the maxillary palps for recording, and, as mentioned above, we consistently observed that 1-octen-3-ol was detected by the neuron with the smallest spikes. However, we cannot rule out the possibility that there are systematic differences in the experimental preparations for SSR between different studies such that different studies target sensilla in different regions on the maxillary palp and that the sensilla in these regions differ in their composition. Future studies analyzing the morphological and molecular composition of sensilla in different parts of the maxillary palp will help in examining this possibility.

The plasticity we observed in LFP and in cpC firing rate appears to be longer lasting (at least 20 s) than the fast adaptation commonly observed in olfactory sensory neurons in insects (Jafari and Alenius 2021; Wicher and Miazzi 2021). Neurons expressing different types of receptors can have different response dynamics (Getahun et al. 2012). Most sensory neurons show fast adaptation, such that the response to even a 1-2 s stimulus rapidly reduces in magnitude during the presence of the stimulus (Nagel and Wilson 2011). The fast adaptation is thought to be downstream of the odor-binding process and at the level of transduction channels, and is likely mediated by changes in calcium ion concentrations (Störtkuhl et al. 1999; Deshpande et al. 2000; Nagel and Wilson 2011). The plasticity in cpC was observed for most odors that activated the neuron; this effect was not observed in cpA and cpB neurons, showing that cpC differs physiologically from the other two neurons. This difference has implications for repeated exposures to complex stimuli involving multiple odorants: for example, as the host odor involves some odorants detected by cpA (such as CO_2_) and some detected by cpC (such as 1-octen-3-ol), the relative ratio of cpA:cpC firing is expected to increase as cpC firing reduces over repeated exposures to the odor.

The ratios of firing rates of different neurons within a same sensillum become even more important when there are inhibitory interactions among the neurons. We indeed observed lateral inhibition between neurons in the *Aedes* capitate peg sensillum. These interactions are likely mediated by ephaptic coupling (Su et al. 2012). We confirmed lateral inhibition from cpC to cpA and cpC to cpB. It is likely that inhibitory interactions are also present among other pairs of neurons, although we could not verify them because of technical limitations. We could not comment on the effect of cpA on other neurons because of the difficulty in detecting cpB and cpC spikes in the presence of large cpA spikes. We also could not comment on the effect of cpB on other neurons because 4-methylcyclohexanol, which activated cpB, also generated some activity in cpC, and thus it would be difficult to ascribe any inhibition triggered by 4-methylcyclohexanol stimulation to cpB firing alone.

In some insects, such as the Japanese beetle *Popillia japonica* and the moth *Helicoverpa zea*, two neighboring neurons within a sensillum detect two molecular components of a pheromone; the exact ratio in which the two neurons are activated allows the insect to discriminate a mating partner from another related insect whose pheromone has a slightly different ratio of the same two molecular components (Cossé et al. 1998; Nikonov and Leal 2002). *Aedes aegypti* has evolved to prefer humans over other animals and this differentiation appears to be based on differences in the relative proportions of molecules in the host odors (Verhulst et al. 2018; Zhao et al. 2022). Notably, the concentration of 1-octen-3-ol in human odor is lower than in cattle odor (Majeed et al. 2016). Different host odors will generate different ratios of activities in the three neurons in the capitate peg sensillum; we speculate that these differences are further enhanced by inhibitory interactions between neurons within the sensillum and could contribute to the anthropophilic behavior in mosquitoes.

## Methods

### Insects

*Aedes aegypti* wild-type (Linnaeus) Liverpool strain, Orco^16/16^ mutants (DeGennaro et al. 2013), and Gr3^4/4^ mutants (McMeniman et al. 2014) were used in this study. Mosquitoes were cultured at 25-28°C with 50-80% relative humidity under a 14 h light: 10 h dark cycle. Larvae were reared in distilled water with fish food (TetraBits). Adult female mosquitoes used for single sensillum recordings were mated but not blood-fed, 4–10 days old, and fed on 10% sucrose solution.

### Single sensillum recordings

Female mosquitoes were anesthetized on ice for 30 s, and then pasted on a mounting setup made with two glass slides. One slide was kept elevated at 30° angle over the other (horizontal) slide using clay. The mosquito’s head was pasted on the elevated slide and the body on the horizontal slide using double-sided tape. To further immobilize the mosquito, epoxy glue was applied on the sides of its eyes and the neck, above the proboscis, the abdomen, and the wings, and below the thorax, the palps, and the antennae. A palp was pasted on the epoxy after rotating it away from the proboscis such that the ventral side, where the capitate peg sensilla are mostly located, faced up. Recording electrodes with tips smaller than 1 μm and reference electrodes with 3-4 μm tips were made from glass capillaries (BF-150-86-10) using P-1000 puller (Sutter), and were filled with physiological saline (pH 7.4, osm 300). Chlorided silver wire inserted in the glass capillary conducted the signal to a pre-amplifier, from where the signal went to a Multiclamp 700B amplifier (Molecular Devices) and then was recorded at the sampling rate of 20,000 samples/s using Digidata 1550A digitizer (Molecular Devices). The preparation was observed using a 20X large-working distance objective on a fixed-stage light microscope (Olympus BX51W1). The recording electrode was put in the center of a capitate peg sensillum while the reference electrode was inserted into a compound eye. The movement of the recording electrode was controlled using a motorized manipulator (Scientifica PatchStar). The odor outlet tube was placed at a distance of 1.5 cm from the animal.

### Odors

Odors used in this study include components of human emanations: 1-octen-3-ol (1OCT3OL), 2-butanone (2BUTONE), 3-methylcyclopentanone (3MCYPONE) (Gallagher et al. 2008), 6-methyl-5-hepten-2-one (6MHO), carbon dioxide (CO_2_), dimethyl trisulfide (DM TS), L-lactic Acid (L-LA), nonanal (NONANAL); plant-derived molecules: (S)-limonene (S-LIMENE), cyclopentanone (CYPTONE) (fruit odor (Horvat and Senter 1984)), geranyl acetate (GER ATE), 1-hexen-3-ol (1-HEXEN-3-OL), beta citronellol (B-CITNOL), linalool (LINALOOL); one plant-derived molecule that also acts as an oviposition attractant for some mosquito species (Bentley et al. 1982): 4-methylcyclohexanol (4MCYHOL); one synthetic repellent: N,N-diethyl-meta-toluamide (DEET); and two solvent controls: mineral oil (MIN OL), and distilled water. All odors were diluted (v/v) in mineral oil except L-Lactic acid, which was diluted in deionized water, and were placed in 50-ml glass bottles. There was a further dilution by a factor of 10 during odor delivery as described below. The concentrations indicated in the Results section reflect the final dilution.

### Odor delivery

The mosquitoes were exposed to a constant stream of dehumidified and filtered compressed air at a flow rate of 2L/min, which was maintained using a 0-3 L/min flowmeter (CVG-Glass). The 2L/min flow was split into two segments that were eventually combined near the animal. One of the two streams was clean air with a flow rate of 1.8 L/min. The other stream with a flow rate of 0.2 L/min, maintained using a 0-0.5 L/min flowmeter, consisted of clean air which passed through an empty bottle during the background period or through the headspace of an odor bottle during the odor delivery period. The switching was done using a three-port solenoid valve (Product code: 11-13-3-BV-24F88, Parker Hannifin) which was controlled by a motor driver (L298N Dual Motor Controller). To check the final odor delivery profile of odors, we used a photo-ionization detector (200B miniPID, Aurora Scientific).

CO_2_ was taken from a compressed cylinder. During the odor period, a stream of CO_2_ with a flow rate of 20 ml/min, maintained using a 1.9-22.8 ml/min CO_2_ flowmeter (Aalborg), was mixed with approx. 1.98 L/min clean air close to the animal, resulting in a final CO_2_ concentration of 0.01.

### Local field potential

LFP was obtained by taking the low-frequency signal from an SSR recording, by applying a 40-Hz low-pass filter in MATLAB. Odor stimulation usually resulted in a negative deflection in LFP. The LFP response to an odor (delivered during 2-3 s interval) was quantified by calculating the most negative value of the LFP in the 2.1-3.1 s (assuming a 0.1 s delay in odor delivery) and subtracting the background LFP value in the two 2 s interval before the start of the odor delivery.

### Spike detection

We use the second derivative of the signal to detect A, B, and C spikes in the SSR recordings from the capitate peg sensillum. The second derivative was calculated in this manner: we first filtered the recording using a 500 Hz low-pass filter. The signal was passed through ‘diff’ function in MATLAB to take the derivative, which was again passed through ‘diff’ to obtain the second derivative. The sign of the peaks is inverted in the second derivatives, so we multiplied the signal by −1 to make the peaks positive. Thresholds were manually selected to separate spikes in A from B, B from C, and C from the background noise. The last of these thresholds was often the most difficult to choose, as the sizes of the C spikes were very small. In most cases, we observed that a small increase or decrease in the threshold did not affect our conclusions: a small change in the threshold affects the spontaneous and the odor-evoked spikes, both, but does not change the fact that the odor-evoked spiking rate is higher than the background when the cell is responding to the odor (or that the firing rates during the background and the odor stimulation period are similar when the cell is not responding to the odor). In some recordings, we also noticed that the sizes of cpA or cpC spikes changed during an ongoing response, particularly when the neurons were firing at high rates.

### Analysis of experiments with 1-octen-3-ol stimulus on top of CO_2_ stimulus

These experiments were performed with trials of duration 20 s. The CO_2_ pulse was given for 3 s in the interval of 2-5 s, and 1-octen-3-ol pulse was given for 1 s in the interval of 3-4 s (i.e., in the middle of the CO_2_ pulse). The firing rate of cpA increased with the onset of the CO_2_ pulse and then reduced gradually within the 3 s trial due to adaptation - the temporal profile of this reduction resembled an exponential decay. When 1-octen-3-ol pulse was delivered, there was an additional reduction in the cpA firing. The effect of 1-octen-3-ol was determined in the following way. We first calculated the instantaneous firing rate during the 2-5 s interval with 20 ms temporal resolution (smoothened with a Gaussian filter), and calculated the time at which the firing rate achieved the peak (t_p_). Then we fitted an exponential decay curve to the firing rate in the interval of t_p_ to 5 s (but excluded 3-4.5 s interval from it, to ensure that fitting was done in only that period when cpA firing was not affected by 1-octen-3-ol). The decay was modeled by the equation: firing rate = c1 + c2 x exp(-(t-t_p_)/c3), where c1, c2 and c3 are parameters obtained by using the ‘fit’ function in Matlab with “NonLinearLeastSquares” method. We finally estimated the effect of 1-octen-3-ol pulse by calculating the difference between the fit and the actual firing rate in the 3.2-4.2 s interval (compared to the odor delivery interval, this interval was shifted by 0.2 s as the effect of the 1-octen-3-ol was maximal in this period).

## Data availability

All datasets are available from the corresponding author upon request.

## Code availability

Code used for analyses are available from the corresponding author on request.

## Acknowledgements

We thank Arun Shankar for help with mosquito breeding. This work was supported by the DBT/Wellcome Trust India Alliance Fellowship [grant number IA/I/15/2/502091], DST/SERB Swarnajayanti Fellowship [SB/SJF/2021-22/04-C], and SERB Core Research Grant [CRG/2020/004719] awarded to N.G.

## Authors Contributions

Conceptualization: NG and SS

Methodology: SS and NG

Data collection (SSR): SS

Data collection (PN Data): PS and Shefali G

Data curation: SS

Data analysis: SS, Sanket G, Smith G, AA and NG

Writing: SS and NG

