## Supplementary Information for "Plasticity and interactions in the odor responses of maxillary palps neurons in *Aedes aegypti*"

*by*

**Singh et al.**

Supplementary Figures S1-S8

**Supplementary Figure S1**

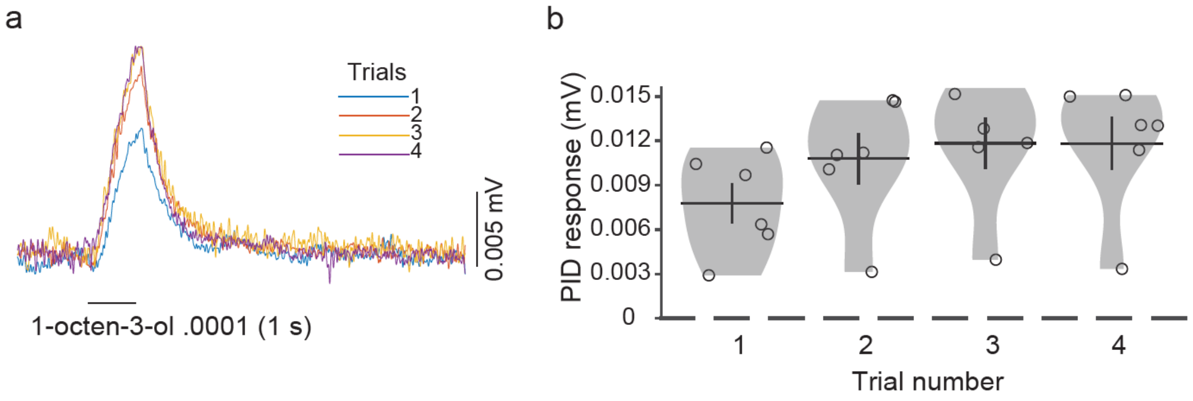


***Figure S1: Odor delivery profile of 1-octen-3-ol .0001***

**a** Representative trace of a photo-ionization detector (PID) measurement showing that the odor (1-octen-3-ol .0001) delivered in the first trial is less than in other trials. Responses were measured by placing the PID nozzle in place of the animal in the direction of odor delivery. **b** Violin plot for the PID responses on 6 different days.

**Supplementary Figure S2**

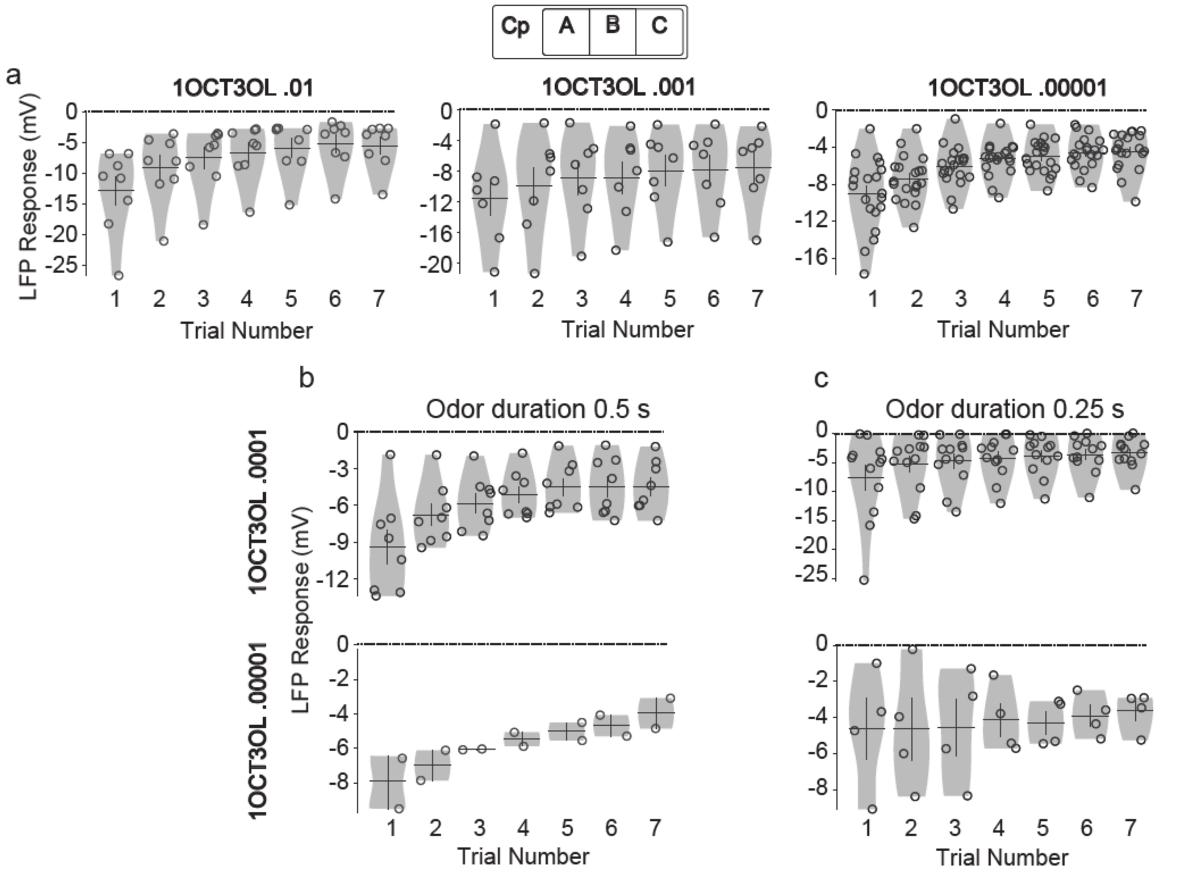


***Figure S2: LFP response plasticity for different concentrations and pulse durations of 1-octen-3-ol***

**a** Violin plots showing reduction over trials in LFP responses in response to 1 s pulses of 1-octen-3-ol .01, 1-octen-3-ol .001, and 1-octen-3-ol .00001. Each point in a violin is from a different sensillum. LFP response plasticity is observed for all three concentrations.

**b,c** Violin plots showing reduction over trials in LFP responses in response to 0.5 s (**b**) and 0.25 s (**c**) pulses of 1-octen-3-ol .0001 and 1-octen-3-ol .00001. At .0001 concentration, plasticity was observed with both pulse durations, while at .00001 concentration plasticity is seen in response to 0.5 s pulse but not with 0.25 s pulse.

**Supplementary Figure S3**

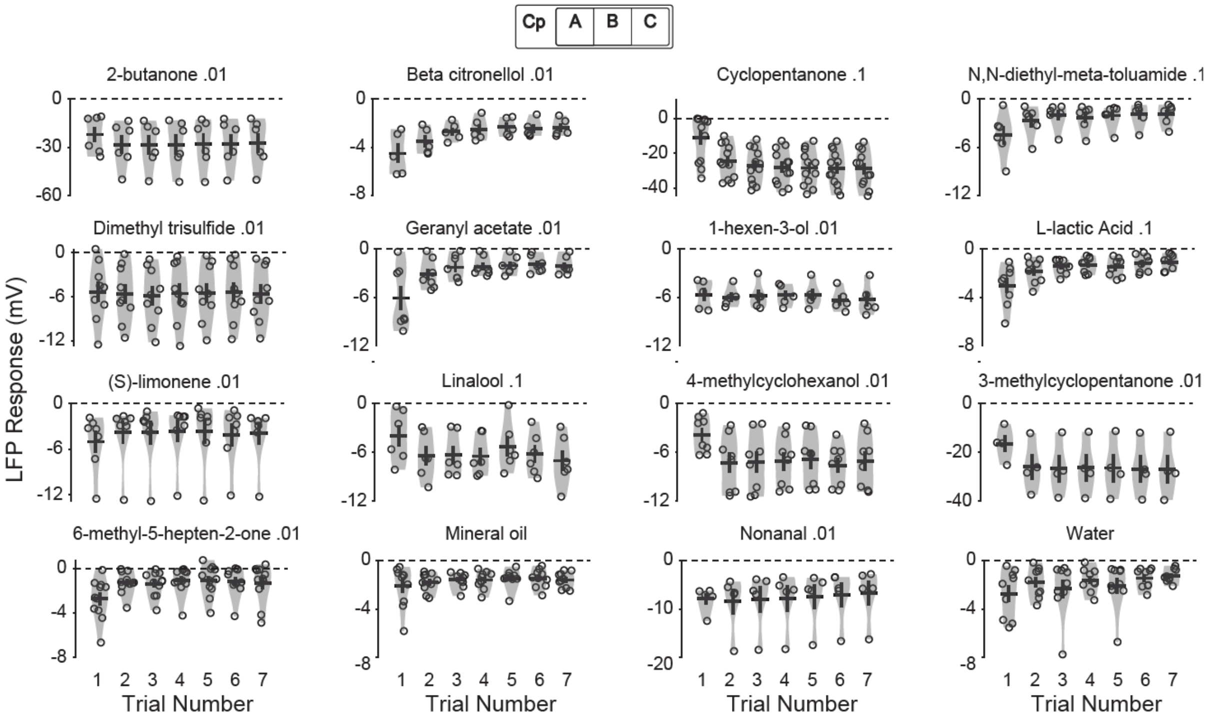


***Figure S3: LFP response plasticity is not unique to 1-octen-3-ol***

Violin plots showing LFP responses in response to 14 odors with 2 solvent controls. Some of the odors tested, including (S)-limonene, β-citronellol, and geranyl acetate show LFP response plasticity with stronger (more negative) response in the first trial and weaker responses in subsequent trials.

**Supplementary Figure S4**

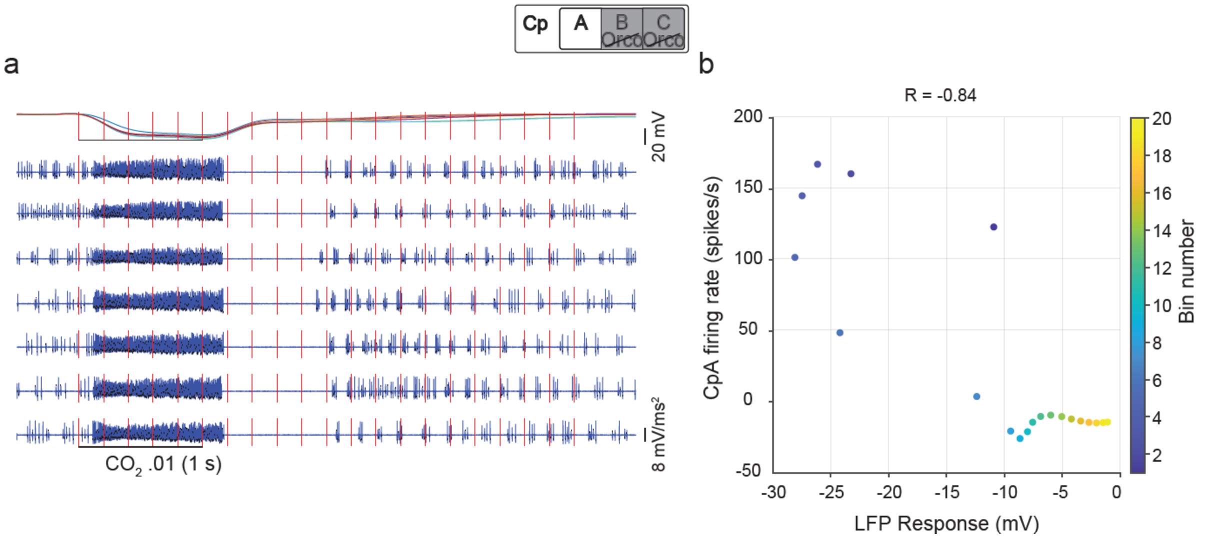


***Figure S4: Correlation between LFP and cpA firing rate***

**a** Representative LFP trace and second derivative traces from an SSR in response to $CO_2$ .01 in an Orco-mutant mosquito. Vertical red lines mark 20 bins of 200-ms duration each starting from odor onset in which the average cpA firing rate responses and average LFP responses were calculated for generating the scatter plot. **b** Scatter plot between the LFP responses and the cpA firing rates over the 20 bins (correlation coefficient is -0.84 in this SSR).

**Supplementary Figure S5**

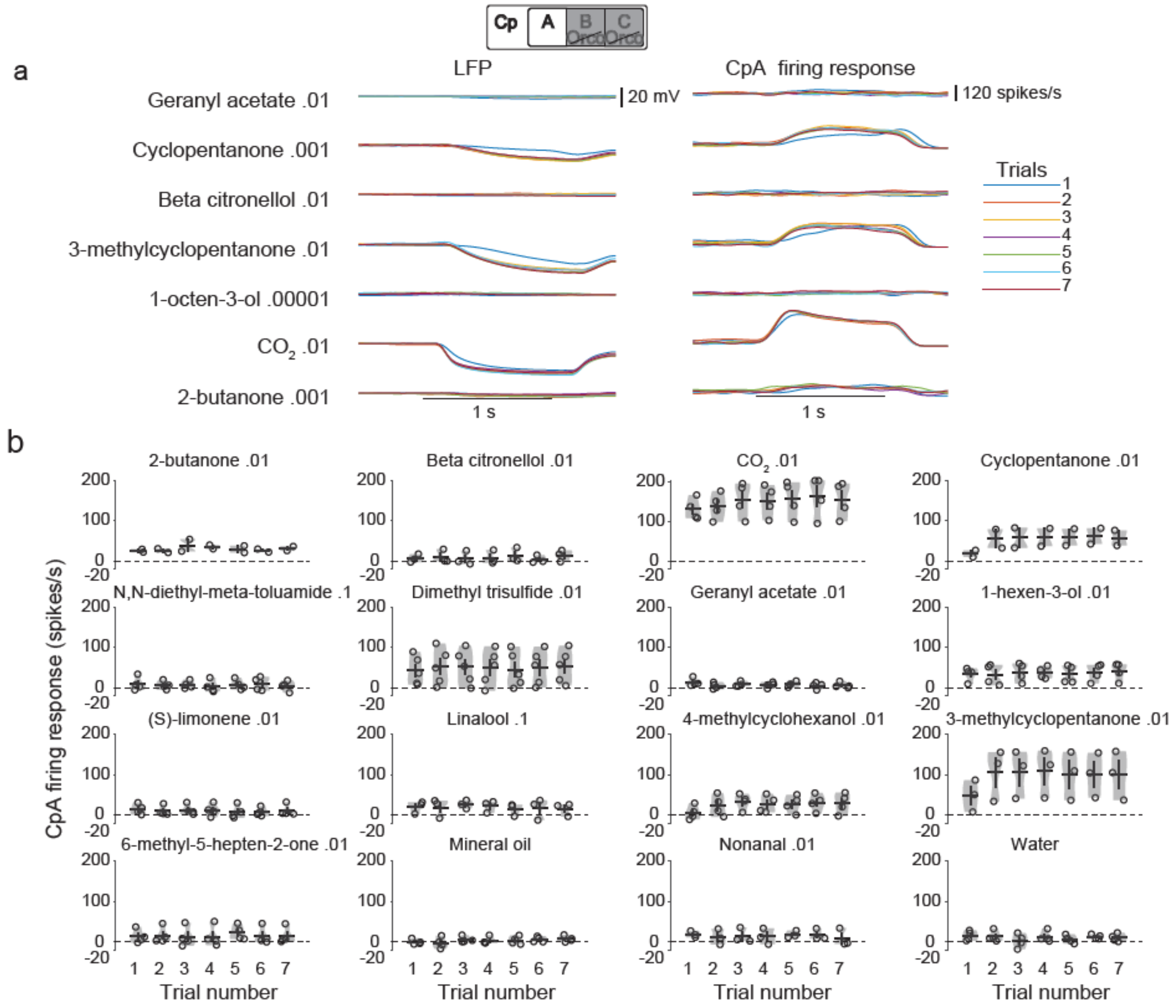


***Figure S5: No plasticity in cpA firing responses***

**a** LFP responses and cpA firing rates in response to 1 s pulses of different odors in Orco-mutant mosquitoes. Each experiment consists of 7 trials (legend on right). No reduction in the response intensity was observed (on the contrary, the response in the first trial was weaker for some odors due to lower odor delivery in the first trial). **b** Violin plots of cpA firing responses from Orco-mutant mosquitoes do not show a reduction in firing rates over trials.

**Supplementary Figure S6**

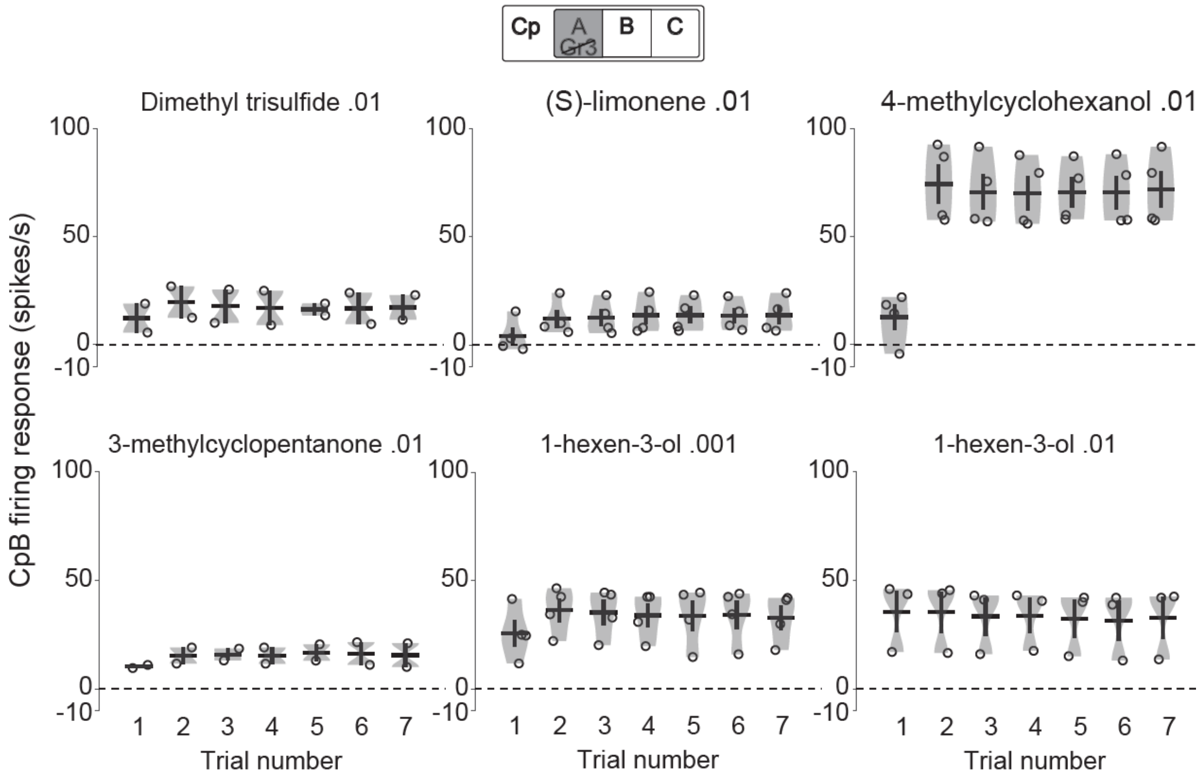


***Figure S6: No plasticity in cpB firing response***

Violin plots show trial-wise cpB firing rates for odors that activate cpB neurons: dimethyl trisulfide, (S)-limonene, 4-methyl-cyclohexanol, 3-methyl-cyclopentanone and two different concentrations of 1-hexen-3-ol. The lower cpB firing response for 4-methyl-cyclohexanol in the first trial is due to lower odor delivery in the first trial.

**Supplementary Figure S7**

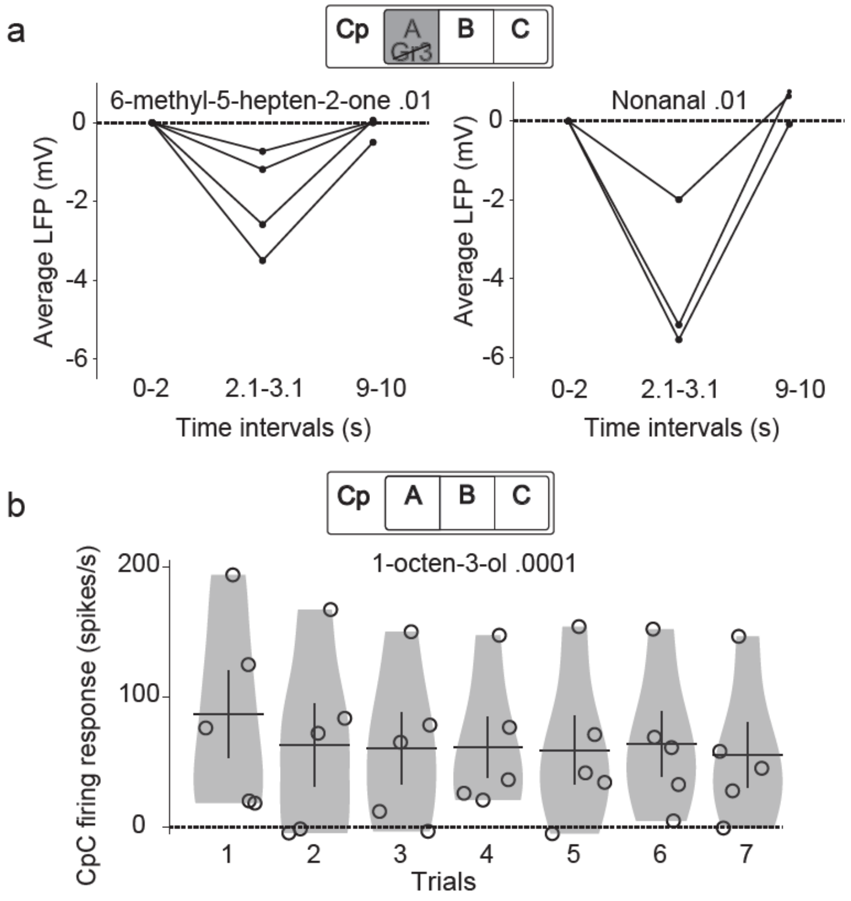


***Figure S7: LFP responses not returning to baseline is not the major reason for plasticity***

**a** Comparing LFP responses to 6-methyl-5-hepten-2-one and nonanal in 3 time-intervals during the 10 s trial: before odor onset (0-2 s), during odor (2.1-3.1 s), and at the end of the trial (9-10 s). The mean values of LFP in the 0-2 s and 9-10 s intervals were not different for 6-methyl-5-hepten-2-one .01 (P=1, n=4, sign-rank test) or nonanal .01 (P=0.5, n=3, sign-rank test). **b** Violin plots show plasticity in cpC firing response to 1-octen-3-ol .0001 in experiments with 20 s trials.

**Supplementary Figure S8**

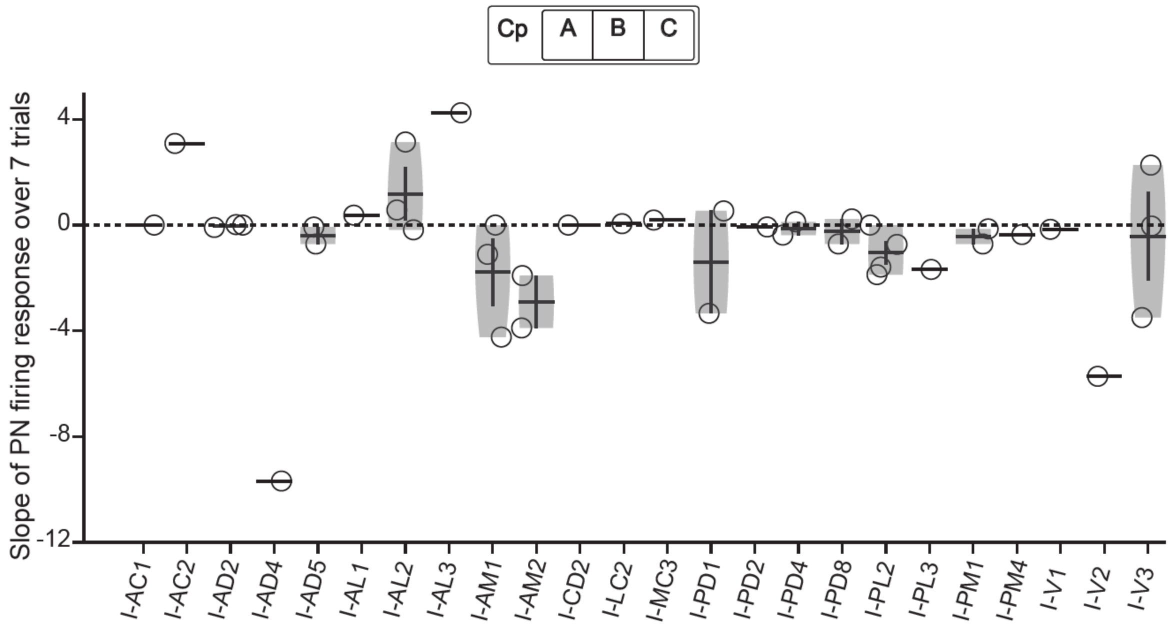


***Figure S8: Comparing plasticity in the responses of different PNs to 1-octen-3-ol .01***

Violin plots showing plasticity in the 1-octen-3-ol responses of 24 different types of PNs. Each point on a violin represents the slope of the fitted line for PN firing response across the 7 trials (a negative slope indicates plasticity). Only those PNs that responded to 1-octen-3-ol were included. Note that most PNs did not have negative slopes (negative slopes observed in a few cases were not consistent across multiple PNs and could be due to chance).
